# Cell atlas of human uterus

**DOI:** 10.1101/267849

**Authors:** Bingbing Wu, Yu Li, Yanshan Liu, Kaixiu Jin, Kun Zhao, Chengrui An, Qikai Li, Lin Gong, Wei Zhao, Jinghui Hu, Jianhua Qian, HongWei Ouyang, XiaoHui Zou

## Abstract

The human uterus is a highly dynamic tissue that undergoes repeated damage repair and regeneration during the menstrual cycle, which make it ideal model to study tissue regeneration and pathological process. Stem/progenitors were speculated to be involved in the regeneration of endometrial epithelial and pathogenesis of endometriosis. But the identity, microenvironment and regulatory mechanisms of the uterus epithelial stem/progenitors in vivo remain unclear. Here, we dissected the cell heterogeneities of the full-thickness human uterus epithelial cells (11 clusters), stroma cells (6 clusters), endothelial cells (5 clusters), smooth muscle cells (2 clusters), myofibroblasts (2 clusters) and immune cells (6 clusters) from 2735 single cell by single cell RNA-seq. Further analysis identified a unique ciliated epithelial cell cluster showing characteristics of stem/progenitors with properties of epithelial-mesenchymal transition (EMT) that mainly localized in the upper functionalis of the endometrium. Ordering the cell subpopulations along the pseudo-space revealed cell clusters possess cellular states of stress, inflammation and apoptosis in the upper functionalis cellular ecosystem of the endometrium. Connectivity map between the human uterus subpopulations revealed potential inflammatory (cytokines and chemokines) and developmental (WNT, FGF, VEGF) signals within the upper functionalis cellular ecosystem of the endometrium, especially from other epithelial clusters, regulating cell plasticity of the EMT-epithelial clusters. This study reconstructed the heterogeneities, space-specific distribution and connectivity map of human uterus atlas, which would provide insight in the regeneration of uterus endometria and reference for the pathogenesis of uterus.

## INTRODUCTION

The human uterus comprises of endometrium, myometrium and the blood vessels. It exhibits remarkable plasticity, the endometria would undergo repeated damage and regeneration, the myometrium would enlarge during pregnancy and return to normal size after gravidity(Ono et al. 2007). Its highly dynamic properties of repeated injury and scar-less repair along the menstrual cycle make it ideal model to study tissue regeneration and pathological process(Maybin and Critchley 2015).

After the proliferative period of the menstrual cycle, the secretory endometria would have two destinies: either to prepare for implantation by fertilized embryos or to shed (Gargett, Nguyen, and Ye 2012; Cakmak and Taylor 2011). The receptive state of the secretory endometria is vital for the proper implantation of embryo, and abnormal physiological conditions would cause miscarriage(Cakmak and Taylor 2011). On the other hand, the shedding endometrial pellets of the secretory endometria would lead to serious pathological outcomes under certain circumstances, like endometriosis, when the shedding endometrial pellets go reversely along the oviduct into the abdominal cavity(Hufnagel et al. 2015). However, the precise regulatory mechanisms for the preparation of the secretory uterus for embryo implantation and etiology of endometriosis remains unclear(Hufnagel et al. 2015).

Endometrial stem/progenitor cells were shown to be involved in the regeneration of the damaged functional layer during the menstrual cycle (Gargett, Nguyen, and Ye 2012) and continuous growth upon embryo implantation, as well as involved in the pathological process of endometriosis(Hufnagel et al. 2015). However, the precise identify, function and regulatory mechanisms of the uterus epithelial stem/progenitors during these physiological and pathological circumstances in vivo remains unclear(Valentijn et al. 2013; Gargett, Schwab, and Deane 2016).

Tissue microenvironment was shown to be indispensable during the tissue development(Camp et al. 2017), homeostasis(Zepp et al. 2017), regeneration and pathological progression(Puram et al. 2017). Single cell analysis has been increasingly utilized to dissect cell heterogeneity and study the dynamic microenvironment between cell subpopulations during biological process such as development, cancer metastasis, tissue homeostasis and pathology(Camp et al. 2017; Puram et al. 2017; Zepp et al. 2017). Thus, in this study, we reconstructed cell atlas of the secretory phase of human uterus through dissecting the cell heterogeneities, reconstructing spatial distribution and connectivity map of the cell clusters by single cell RNA-seq.

## RESULTS

### Single cell RNA-seq of full-thickness secretory human uterus

First, we used drop-seq (10x Genomics) based single cell RNA-seq to profile single cell suspension from a full-thickness secretory human uterus tissue (Fig 1A). Then we used the Cell Ranger Pipeline (10x Genomics) to analyze the unique molecular tagged (UMI) Of the raw sequencing data (Fig S1A). We profiled 2735 individual cell of the human uterus (Fig S1B). We obtained saturated sequencing with about 680k reads per cell (Fig S1C), collected 2735 cells with high UMI counts, and the median gene number detected per cell was about 3,183 (Fig S1D).

**Figure 1.**
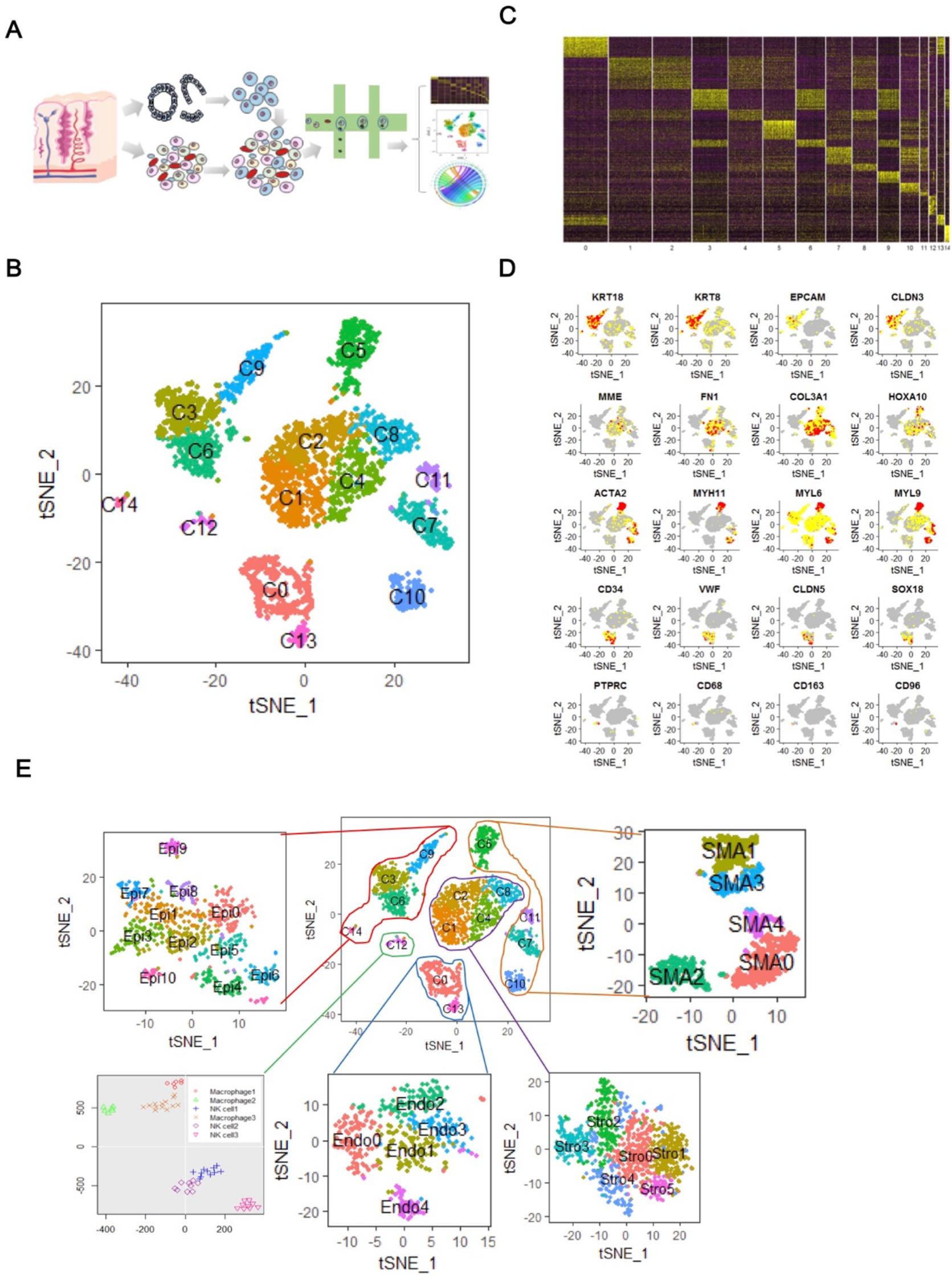
Single cell RNA-seq of full-thickness human uterus. (A) Workflow shows sample processing, enzymatic digestion and drop-seq based single cell RNA-seq. (B) t-distributed stochastic neighbor embedding (t-SNE) plot of 2735 single cell from a full-thickness secretory phase uterus by single cell RNA-seq (scRNA-seq). (C) Heatmap shows differential expressed gene signatures of each cell cluster from scRNA-seq. (D) t-SNE plot of selected marker genes from gene signatures of each cell cluster (KRT18, KRT8, EPCAM, CLDN3 for epithelial cell; MME, FN1, COL3A1, HOXA10 for stroma cells; ACTA2, MYH11, MYL6, MYL9 for SMA+ cells (smooth muscle cells and myofibroblasts); CD34, VWF, CLDN5, SOX18 for endothelial cells; PTPRC, CD68, CD163, CD96 for immune cells). (E)Each cell cluster (epithelial cell, stroma cell, endothelial cell, SMA+ cell, immune cells) was further clustered into their relevant subpopulations (32 sub-clusters in total).

Unsupervised clustering based on principal components of the most variable expressed genes (Satija et al. 2015) partitioned all the cells into 15 clusters, which we visualized using t-distributed stochastic neighbourhood embedding (t-SNE) (Fig 1B) and principal component analysis (PCA) (Fig S1E, S1F), each cluster possesses a unique set of signature genes (Fig 1C). As labeled by known marker genes, there are clusters of endometrial epithelia, stroma, endothelia, SMA+, immune cells in human uterus. Endometrial epithelial cells express high level of KRT8, KRT18, EPCAM and CLDN3, endometrial stromal cells express high level of MME, FN1, COL3A1, HOXA10, endothelial cell express high level of CD34, VWF, CLDN5, SOX18, SMA+ cells express high level of ACTA2, MYH11, MYL6, MYL9. Endometrial immune cell express high level of PTPRC, CD68, CD163 and CD96 (Fig 1D).

To further investigate cellular heterogeneity of each cluster of the full-thickness human uterus, we cluster the endometrial epithelia, stroma, endothelia, SMA+, immune cells separately, and we acquired 11 sub-clusters of endometrial epithelial cells, 5 sub-clusters of endmotrial stromal cells, 5 sub-clusters of SMA+ cells, 5 sub-clusters of endothelial cells, 3 sub-clusters of macrophages and 3 sub-clusters of natural killer cells (Fig 1E). Totally, we found 32 sub-clusters from 5 main groups.

### Heterogeneity of uterus epithelia cells

Analysis of each unique signature genes of endometrial epithelial sub-clusters (Fig 2A, 2B) showed heterogeneity of ciliated and secretory epithelial sub-clusters. There are 5 ciliated epithelial sub-clusters (Epi4, Epi5, Epi6, Epi9, Epi10) as labelled by ciliated marker alpha-tubulin expression (TUBA1A and TUBA1B) (Fig 2C). There are 7 secretory epithelial sub-clusters (Epi0, Epi1, Epi2, Epi3, Epi7, Epi8, Epi9) in the uterus as labelled by secretory marker secretoglobin family (SCGB1D4, SCGB2A1) and inflammatory cytokines and chemokines (CXCL8, VEGFA) (Fig 2D). There is a sub-cluster of cells (Epi9) that express both the ciliated epithelial marker (TUBA1A) and secretory epithelial markers (SCGB1D4, SCGB2A1) (Fig 2C,2D).

**Figure 2.**
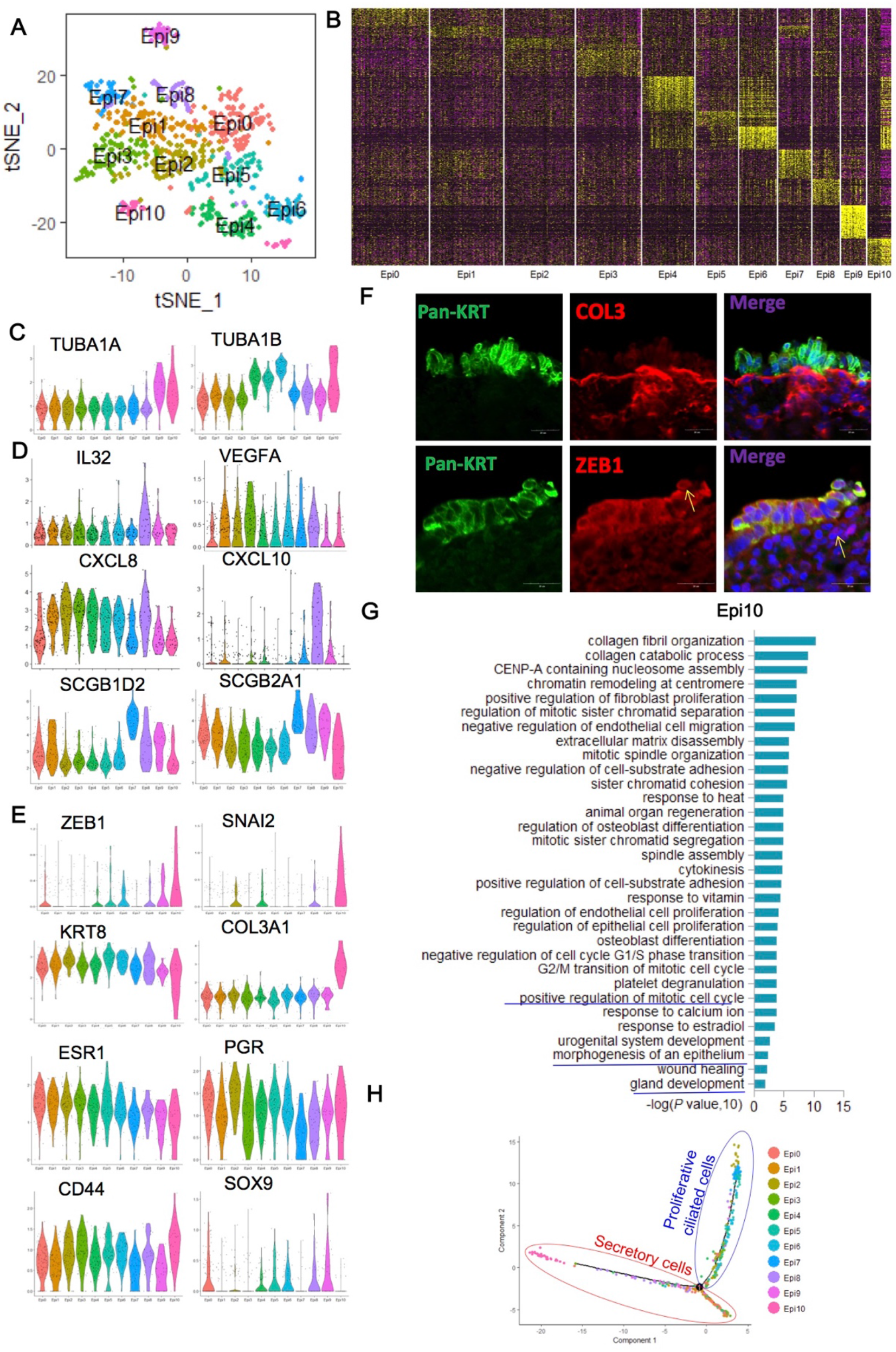
Heterogeneity of uterus epithelia subsets. (A) t-distributed stochastic neighbor embedding (t-SNE) plot of epithelial cells using Seurat. (B) Heatmap shows differential expressed gene signature of each sub-cluster from epithelial cells. (C) Violin plot depict markers (TUBA1A and TUBA1B) of ciliated epithelial sub-clusters (Epi4, Epi5, Epi6, Epi9, Epi10). (D) Violin plot depict markers of secretory epithelial sub-clusters (Epi0, Epi1, Epi2, Epi3, Epi7, Epi8, Epi9) as labelled by secretoglobin family (SCGB1D4, SCGB2A1) and inflammatory cytokines and chemokines (CXCL, VEGFA). (E) Violin plot depict markers (KRT8, COL3A1) and transcription factors (ZEB1, SNAI2) of EMT epithelial sub-cluster(Epi10). (F) Immunofluorescent staining showed the expression of EMT markers (pan-KRT, COL3A1, ZEB1) in the upper layer of the functionalis of the endometria. (G) Gene ontology analysis of the highly expressed genes in the EMT sub-cluster (Epi10). (H) Pseudospace ordering of all the epithelial subclusters.

Interestingly, we found a sub-cluster of ciliated epithelial cells (Epi10) that possess epithelial-mesenchymal transition (EMT) characteristics, which express both epithelial markers (KRT8) and stroma cell markers (COL3A1), as well as EMT transcription factors (SNAI2, ZEB1) (Fig 2E), which were further confirmed by immunofluorescent staining that the EMT epithelial cells were mainly localized in the upper layer of the functionalis of the endometria (Fig 2F). This EMT sub-cluster express low level of estrogen receptor (ESR1) and high level of CD44 (Fig 2E) (a previous reported mouse endometrial stem cell marker(Janzen et al. 2013)). Gene ontology (GO) analysis of the highly expressed genes in this EMT sub-cluster showed enrichment of GO terms associated with morphogenesis and development of epithelia and gland (Fig 2G). All this imply the uniqueness and association of this EMT sub-cluster with endometrial epithelial stem/progenitor cells.

In order to reconstruct the spatial distribution of the epithelial sub-clusters, we conducted monocle (Satija et al. 2015) to align the 11 subpopulations along the pseudo-space (Fig S2A). Results showed that cluster Epi10, Epi6, Epi4 and Epi5 distributed at the beginning of the pseudo-space, while cluster Epi8, Epi7 and Epi9 distributed at the end of the pseudo-space (Fig S2B). GO analysis with the highly expressed genes of cluster Epi10, Epi6, Epi4 and Epi5 showed enrichment of GO terms of cell cycle and DNA synthesis (Fig 2G, Fig S3C-E). GO analysis with the highly expressed genes of cluster Epi2, Epi3, Epi7 and Epi8 showed enrichment of GO terms of response of hypoxia, regulation of apoptotic process and interferon signaling pathway (Fig S3A, S3B, S3F, S3G). These results indicated a ciliated-secretory distribution pattern of the epithelial cells along the uterus cavity (Fig 2H).

### Heterogeneity of uterus stroma cells

Further clustering the stroma cells of uterus revealed 6 distinct cell populations with their unique molecular signatures (Fig 3A, 3B). In order to reconstruct the spatial distribution of the stroma cell, we conducted monocle to align the 6 subpopulations along the pseudo-space (Fig S4A). Results showed that cluster Stro2, Stro3 and Stro4 distributed at the beginning of the pseudo-space, while cluster Stro0, Stro1 and Stro5 distributed at the end of the pseudo-space (Fig S4B). GO analysis with the highly expressed genes of cluster Stro2 and Stro3 showed enrichment of GO terms of cell adhesion, cell differentiation and regulation of wound healing (Fig 3D, 3E, 3F). These results indicated that the cluster Stro2, Stro3 and Stro4 were corresponding to stroma cells from the basal layer of the endometrial, which was responsible for the cyclic regeneration of the shedded functional layer. GO analysis with the highly expressed genes of cluster Stro0, Stro1 and Stro5 showed enrichment of GO terms of response of hypoxia and regulation of apoptotic process (Fig 3C, 3G). These results implied that the cluster Stro0, Stro1 and Stro5 were corresponding to stroma cells from the functional layer of the endometrial, which was far away from the blood vessels and would undergo apoptosis upon shedding during each menstrual cycle. These results indicated a basal-functional distribution pattern of the stroma cells along the uterus cavity (Fig 3H).

**Figure 3.**
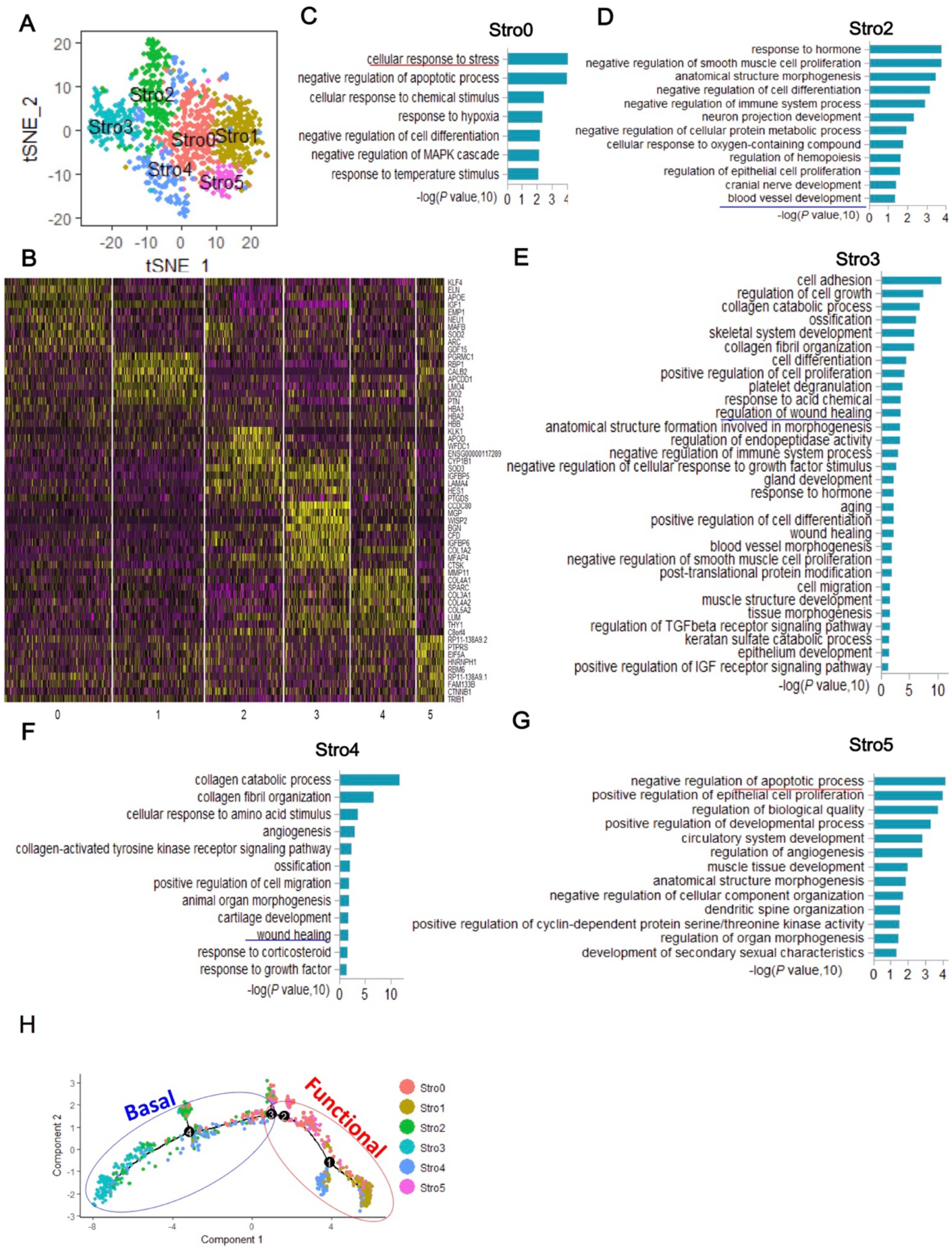
Heterogeneity of uterus stroma subsets. (A) t-SNE plot of stromal cells using Seurat. (B) Heatmap shows differential expressed gene signature of each sub-cluster from stromal cells. (C) Gene ontology analysis of the highly expressed genes in the stromal sub-cluster Stro0. (D) Gene ontology analysis of the highlyexpressed genes in the stromal sub-cluster Stro2. (E) Gene ontology analysis of the highly expressed genes in the stromal sub-cluster Stro3. (F) Gene ontology analysis of the highly expressed genes in the stromal sub-cluster Stro4. (G) Gene ontology analysis of the highly expressed genes in the stromal sub-cluster Stro5.(H) Pseudospace ordering of all the stromal sub-clusters.

### Heterogeneity of uterus endothelia cells

Blood vessels were vital for the regeneration of the uterus endometria, and the endothelia cells would also undergo repeated shedding and regeneration during each menstrual cycle(Maybin and Critchley 2015). Further clustering the endothelia cells of uterus revealed 5 distinct cell populations with their unique molecular signatures (Fig 4A, 4B, 4C). In order to reconstruct the spatial distribution of the endothelia cell, we conducted monocle to align the 5 subpopulations along the pseudo-space (Fig S5A). Results showed that the cluster Endo4, Endo3 and Endo1 distributed at the beginning of the pseudo-space, while the cluster Endo2 and Endo0 located at the end of the pseudo-space (Fig S5B). GO analysis revealed that genes associated with T cell co-stimulation and antigen processing and presentation were highly expressed in cluster Endo2 (Fig 4F). Genes associated with angiogenesis and endothelial cell differentiation were highly expressed in cluster Endo1 and Endo4 (Fig 4E,4G). These results also indicated a basal-functional distribution pattern of the endothelia along the uterus cavity (Fig 4H). As the functional region of the endothelia would shed during the menstrual phage, which was mainly mediate by immune cells, and the basal region of the endothelia would be responsible for the regeneration of the blood vessel in the functional layer.

**Figure 4.**
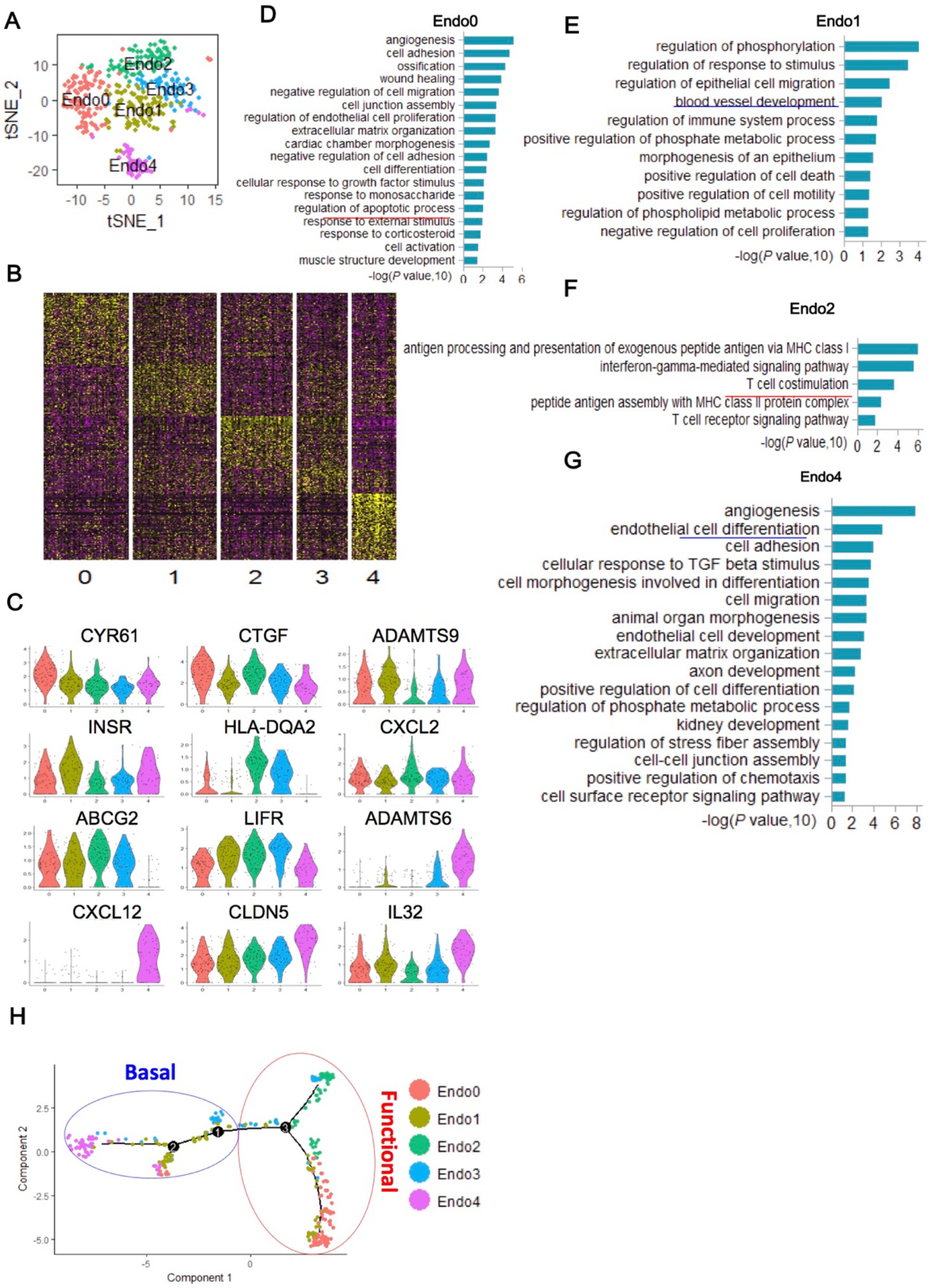
Heterogeneity of uterus endothelial subsets. (A) t-SNE plot of endothelial cells using Seurat. (B) Heatmap shows differential expressed gene signature of each sub-cluster from endothelial cells. (C) Violin plot depict selected markers of each endothelial sub-cluster. (D) Gene ontology analysis of the highly expressed genes in the endothelial sub-cluster Endo0. (E) Gene ontology analysis of the highly expressed genes in the endothelial sub-cluster Endo1. (F) Gene ontology analysis of the highly expressed genes in the endothelial sub-cluster Endo2. (G) Gene ontology analysis of the highly expressed genes in the endothelial sub-cluster Endo4. (H) Pseudospace ordering of all the endothelial sub-clusters.

### Heterogeneity of uterus smooth muscle and myofibroblasts

Further clustering the SMA+ cells of uterus revealed 5 distinct cell populations with their unique molecular signatures (Fig 5A, 5B). We identified 2 clusters of smooth muscles (SMA1 & SMA3), which expressed high level of ACTG2, ACTA2 and DES (Fig 5C). We also found 2 clusters of myofibroblasts (SMA0, SMA4), which expressed high level of COL4A1 and ACTA2 (Fig 5C). Both of the myofibroblasts were in stress (Fig 5D) or inflammatory states (Fig 5G) according to their GO analysis of their highly expressed signature genes. Myofibroblasts were involved in the cyclic menstrual injury and scarless regeneration through wound contraction and synthesis and remodeling of ECM (Maybin and Critchley 2015). Excessive activation of myofibroblasts caused by abnormal regulations were highly correlated with adenomyosis(Ibrahim et al. 2017) and endmetriosis (van Kaam et al. 2008) through abnormal deposition of extracellular matrix from myofibroblast, the discovery of its dysregulation factors may shed light on the pathogenesis of these diseases(Liu et al. 2016).

**Figure 5.**
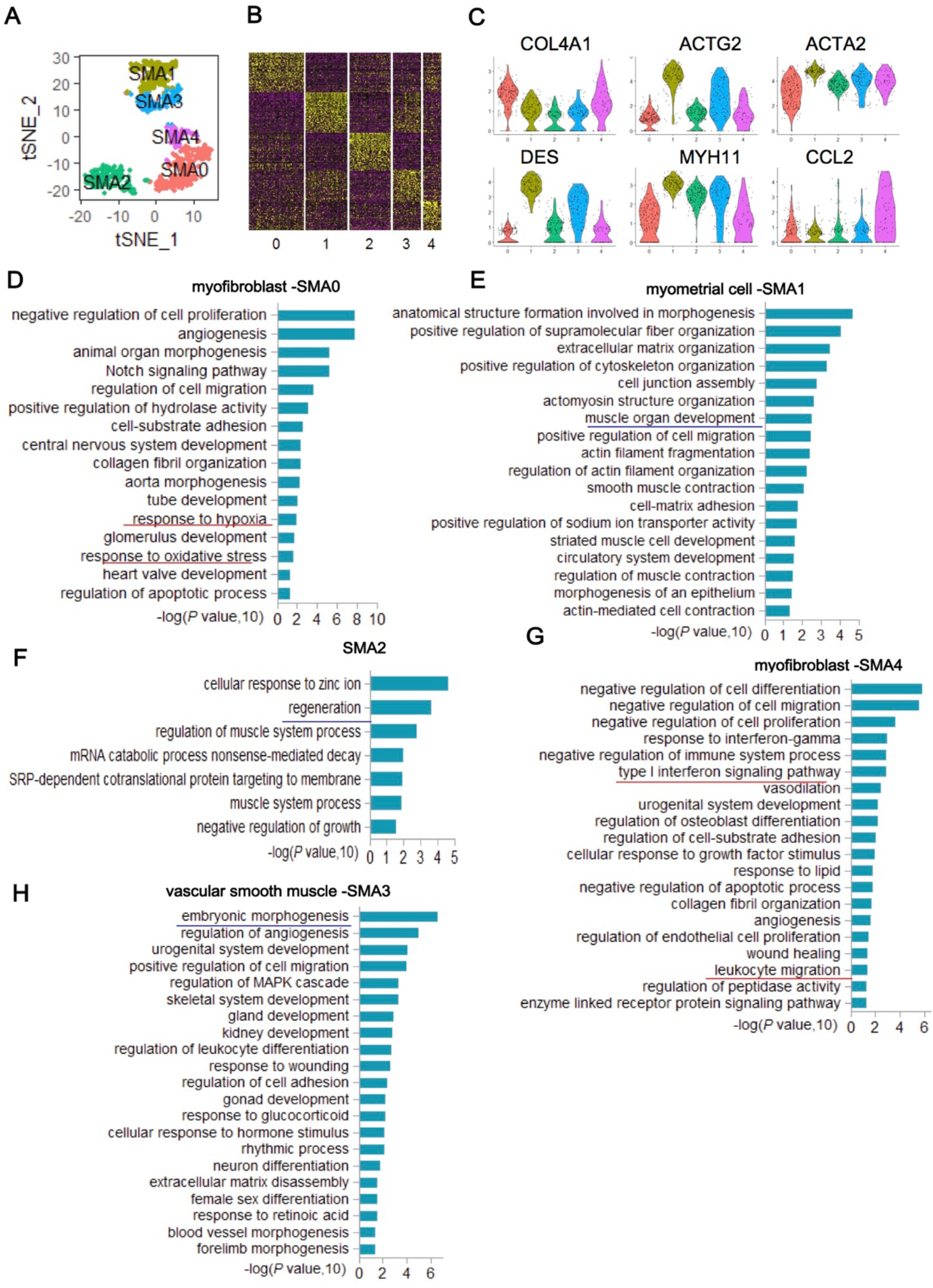
Heterogeneity of uterus smooth muscle cell and myofibroblast subsets. (A) t-SNE plot of SMA+ cells using Seurat. (B) Heatmap shows differential expressed gene signature of each sub-cluster from SMA+ cells. (C) Violin plot depict selected markers of smooth muscle cell (SMA1, SMA3) markers (ACTG2, ACTA2, DES, MYH11) and myofibroblast (SMA0, SMA4) markers (COL4A1, ACTA2). (D) Gene ontology analysis of the highly expressed genes in the myofibroblast SMA0. (E) Gene ontology analysis of the highly expressed genes in the myometrial cell SMA1. (F) Gene ontology analysis of the highly expressed genes in the sub-cluster SMA2. (G) Gene ontology analysis of the highly expressed genes in the myofibroblast SMA4. (H) Gene ontology analysis of the highly expressed genes in the vascular smooth muscle cell SMA3.

### Heterogeneity of uterus immune cells

Immune cells were highly correlated with physiology and pathology of the uterus(Cousins et al. 2016; Fu et al. 2017). Unsupervised clustering of the immune cells in human uterus revealed 3 sub-cluster of macrophages and 3 cub-clusters of natural killer cells (Fig 6A), as labeled by the expression of CD86 and CD96 (Fig 6B), respectively, each sub-cluster of macrophage (Fig 6C) and natural killer cells (Fig 6D) possess their unique molecular signatures. The 3 macrophage clusters showed distinct expression of functional markers (CSF1R and TLR4, Fig 6E). NK cell subsets were reported to rebuild and maintain appropriate local microenvironment for fetal growth during early pregnancy(Fu et al. 2017). Distinct subsets of monocytes/macrophages were spatio-temporally distributed, responsible for breakdown, repair and remodeling, respectively(Cousins et al. 2016). Abnormal interactions between the immune cells and the endometrial tissues were highly correlated with endometriosis, the abnormal NK cell activity would cause inadequate removal of menstrual debris and macrophages would further facilitate the proliferation of the menstrual debris in peritoneal cavity (Seli and Arici 2003).

**Figure 6.**
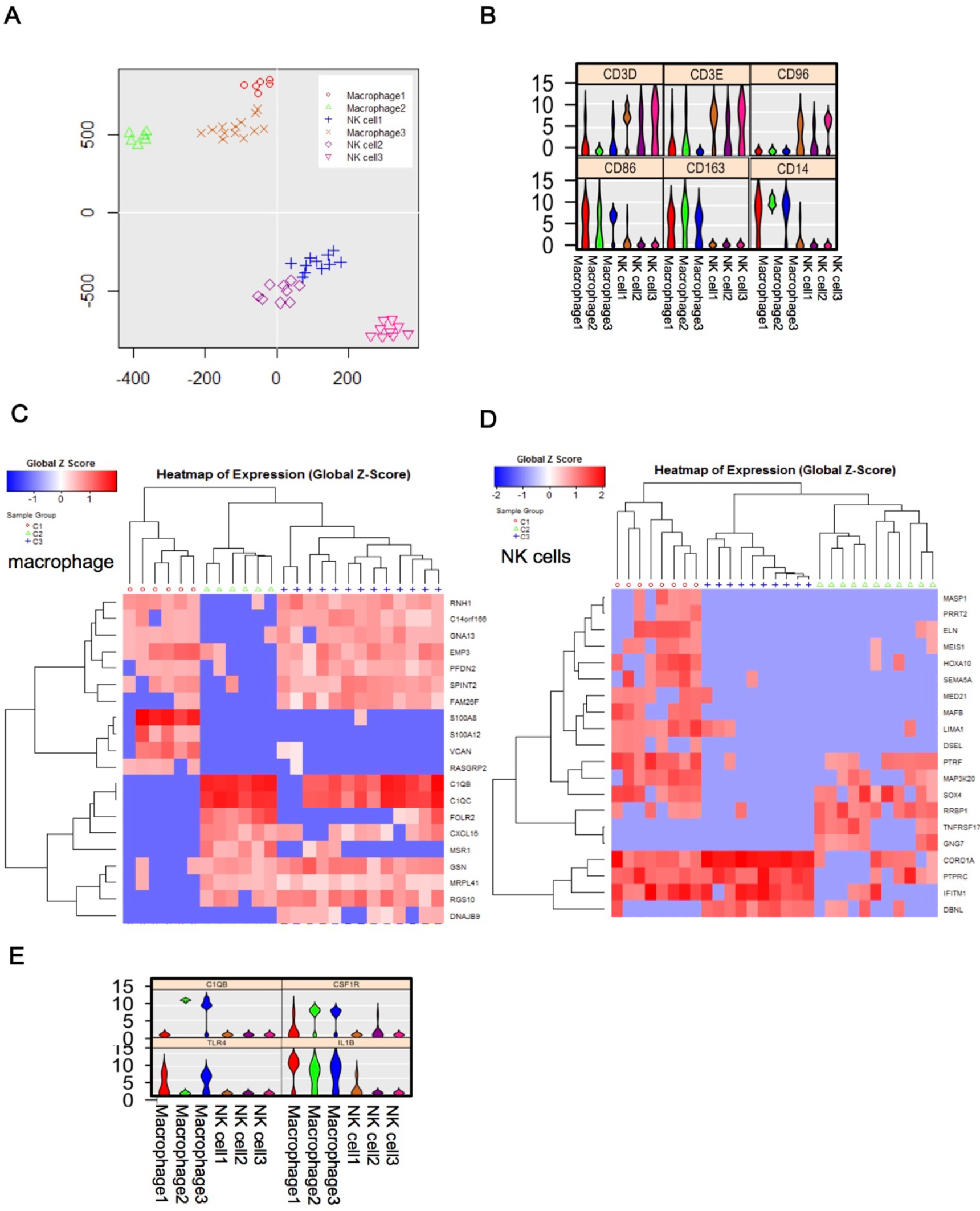
Heterogeneity of uterus immune cell subsets. (A) t-SNE plot of immune cells depict sun-clusters of macrophage and NK cells. (B) Violin plot depict selected markers of macrophage(CD86,CD163) and NK cells(CD96). Heatmaps show Top20 differential expressed gene signature of each sub-cluster from macrophage(C) and NK cells(D). (E) Violin plot depict selected functional markers (C1QB, CSF1R, TLR4,IL1B) of macrophage and NK cells.

### Stress, inflammatory and apoptotic ecosystem of the uterus endometria

The uterus forms unique compartmentalized econsystem of stress, inflammation and apoptotic, which contain epithelial subpopulations (Epi 2, 3, 4, 6, 7, 8, 10), stroma cell subpopulations (Stro0, 1, 5), endothelial cell subpopulations (Endo 0, 2), myofibroblasts (SMA 0, 4) and immune cells (macrophage 1, 2, 3 and NK cell 1, 2, 3), according to the pseudo-space (Fig S4, S5) and GO analysis (Fig S3, 3-5). We next verified these results by using immune fluorescence. The results showed that the upper functionalis layer of the endometria was surrounded by more CD45+ lymphocyte compared with those of the basalis and myometrial layer of the uterus (Fig 7A). While the CD68+ macrophages were evenly distributed in the three layer of the uterus (Fig 7B). We use TUNEL kit to detect the number of apoptotic cells in the full-thickness uterus. There are more TUNEL positive staining in the upper functionalis compared with those of the basalis and myometrial layer of the uterus (Fig 7C). Next we use γH2A.X as marker of DNA damage marker, and we found that there are abundant γH2A.X positive staining in the upper functionalis, even in the some of the epithelial cells, while there’s hardly any γH2A.X positive cells in the basalis and myometrial layer of the uterus (Fig 7E). Our results confirmed that there was a compartmentalized, stress, inflammatory and apoptotic ecosystem in the upper layer of the functionalis of the uterus endometria.

**Figure 7.**
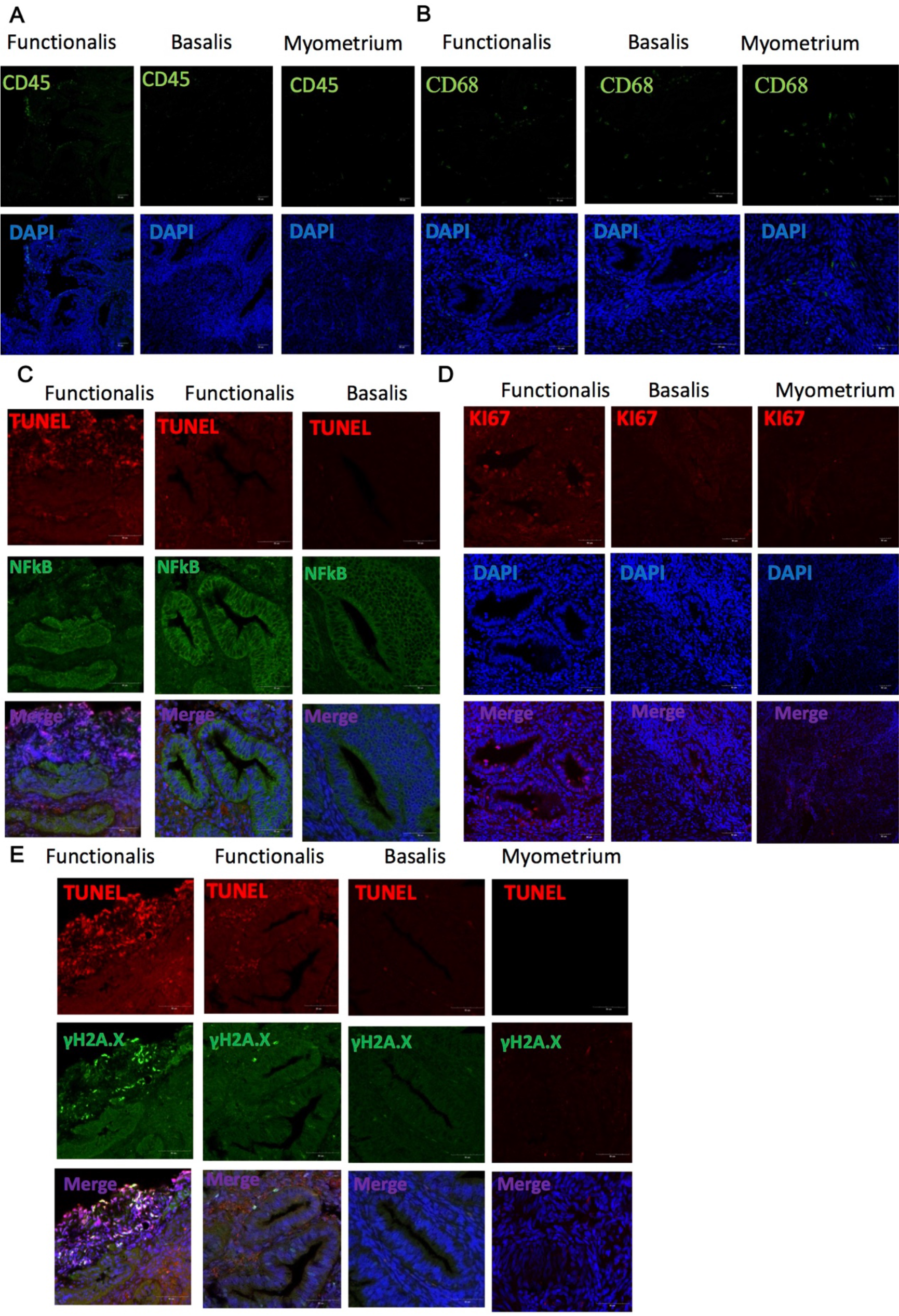
Stress, inflammatory and apoptotic subsets in the uterus ecosystem. Immunofluorescent staining showed the representative number of CD45 positive immune cells (A, green), CD68 macrophages (B, green), TUNEL positive apoptotic cells (C & E, red), NFkB inflammatory state cells (C, green), proliferative cells (D, red) and DNA damage γH2A.X + cells (E, green) in the secretory full-thickness human uterus (three layers: functionalis, basalis layer and myometrium). All the nucleus was stained with DAPI(blue). Scale bar,50 *μ*m.

### Connectivity map of human uterus cells reveal microenvironment regulating epithelial plasticity

As different cell subpopulations were physically surrounded by each other, communications among cells would regulate cell state and even determine cell fate(Camp et al. 2017; Zepp et al. 2017). Thus, we reconstructed the intra-uterus connectivity map among subpopulations by using known ligand-receptor pairs. Finally, we get 1024 connections of 2000 ligand-receptor pairs from 32 sub-clusters (Fig 8A).

**Figure 8.**
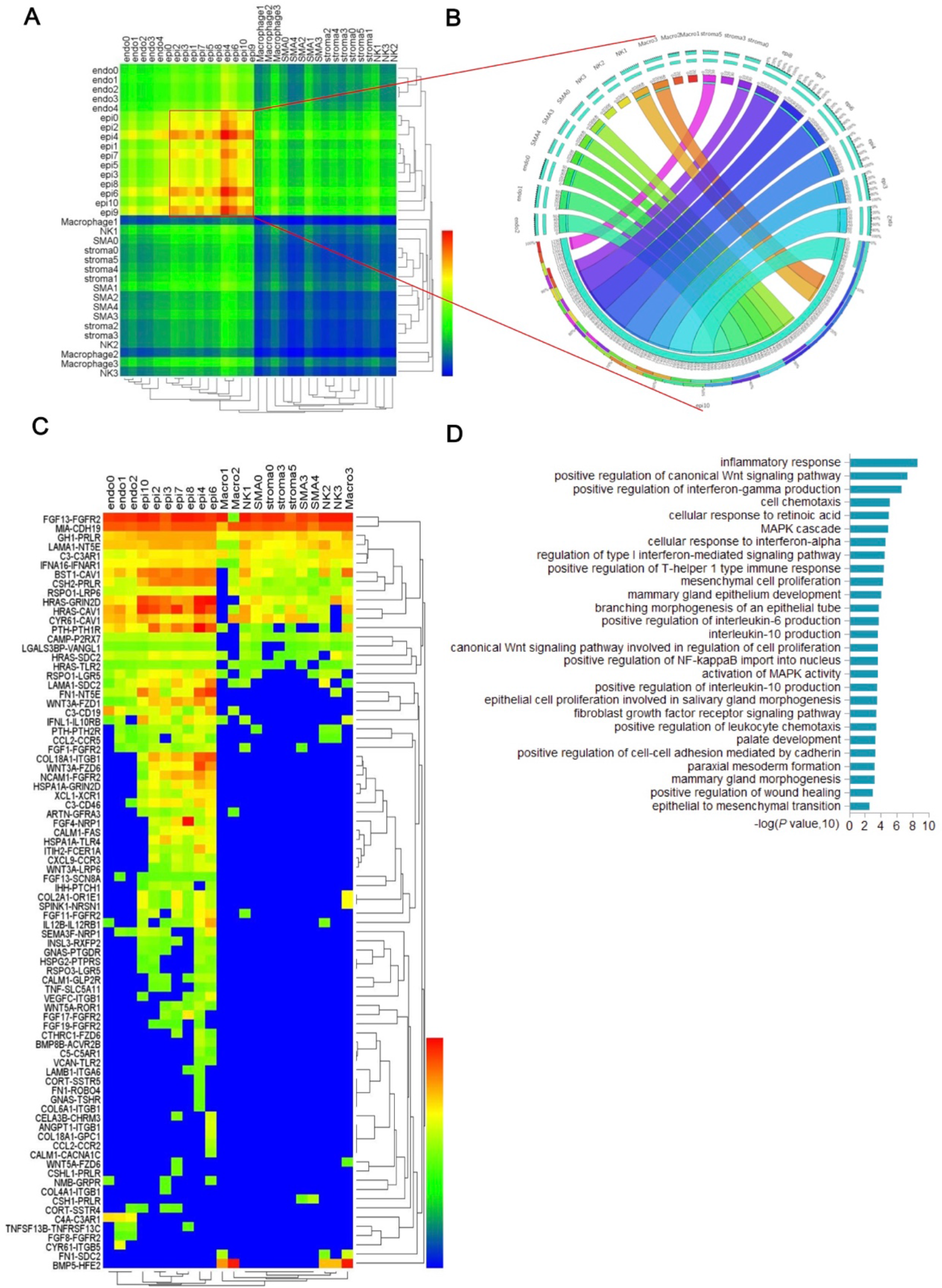
Connectivity map of human uterus subsets. (A) Heatmap shows the total numbers of putative receptor-ligand interactions between two sub-clusters from the ecosystem of the full-thickness secretory uterus. (B) Circle plot depicts the total number of ligand-receptor interactions between each sub-cluster from the EMT microenvironment and the EMT sub-cluster (epi10). (C) Heatmap shows the top ligand-receptor interactions between each sub-cluster from the EMT microenvironment and the EMT sub-cluster. (D) Gene ontology analysis of the ligands from the top ligand-receptor interactions between each sub-cluster from the EMT microenvironment and the EMT sub-cluster.

Nest we studied the potential regulatory microenvironment of the epi10 subpopulation with EMT properties using the connectivity map. As the epi10 cell population localized at the upper layer ecosystem of the functionalis of the uterus (Fig 8B) with the common characteristics of stress, inflammation and apoptotic. Connectivity map of the epi10 revealed that all the epithelial cell subpopulations showed more ligand-receptor pair interactions to epi10 compared to those of the other cell populations (Fig 8B). Further analysis of the ligands secreted into the microenvironment of the epi10 showed abundant extracellular matrix (LAM, FN1, COL), growth factors (FGF, WNT, BMP, VEGF,), inflammatory cytokines (C3, TNF, CXCL, IL12, CCL2) (Fig 8C), which showed enrichment of GO terms of inflammatory response, WNT signaling, epithelial cell proliferation, mesoderm formation and epithelial to mesenchymal transition (Fig 8D). These results implied that the fate of the EMT epithelial cluster was regulated and reprogrammed by its microenvironment, which may reprogram other differentiated epithelial cells into the EMT cluster.

## DISCUSSION

Heterogeneity and cross-talk among subpopulations of complex tissues and organs regulate development. homeostasis, regeneration and pathology(Camp et al. 2017; Zepp et al. 2017; Puram et al. 2017), which remain great challenges until the wide-spread applications of single cell RNA-seq. Here, we reconstructed an atlas of the human uterus tissues: Our atlas provided the most detailed cell diversity of the uterus tissue so far. Our atlas provided a compartmentalized cell ecosystem. Our atlas highlighted a EMT program in the epithelial subset. Our atlas provided a dynamic connectivity map of the uterus with diverse communications in the upper functionalis layer of the endometria regulating the EMT program.

### Our atlas provided the most detailed cell diversity of the uterus tissue

Most of previous study on uterus biology were based on bulk uterus/endometrium tissue transcriptomics analysis(Diaz-Gimeno et al. 2014) or comparison between different region of the tissue(Evans et al. 2014). As the advance of technology and development analysis pipelines, studies have come to the single cell level(Proserpio and Lonnberg 2016). One of our previous study on uterus epithelial development was based on single cell analysis(Wu et al. 2017). A previous report compared the transcriptomics of endometrial stroma cells in vivo and in vitro by single cell RNA-seq (Krjutskov et al. 2016). We found 32 functional distinct sub-clusters from 5 main groups of the full-thickness uterus tissues in total. Our atlas thus provided the most detailed cell diversity of the uterus tissue so far.

### Our atlas provided a compartmentalized cell ecosystem

We discovered subsets of endmotrial epithelial, stromal, endothelial cells and myofibroblasts with distinct states, some subsets showed state of proliferation, while others were in hypoxia, stress and inflammatory states, the rest were in states of development, wound healing and regeneration states. The hypoxia and stress were reported to be functional for the angiogenesis, proliferation and metabolism during the menstrual cycle of the uterus that resemble the process of ischemia and reperfusion (Maybin and Critchley 2015).

### Our atlas highlighted a EMT program in the epithelial subset of the uterus

Our study discovered an unique subset of epithelial cells of the uterus with stem/progenitor property of proliferation, EMT and low hormone receptor expression, which was similar to the characteristics of the epithelial cell cluster we previous reported during the development of mice uterus that was also highly proliferative, EMT and low hormone receptor expression and located in the luminal layer of the uterus (Wu et al. 2017).

Previous reports hypothesized that potential human endometrial epithelial stem/progenitor cells were mainly located in the gland of the basalis layer of the endometrium (Gargett 2007; Nguyen, Sprung, and Gargett 2012). SSEA1 positive cells were shown to possess some stem/progenitor cell properties (longer telomerase activities, more quiescent and lower expresion of estrogen receptor) in vitro, which mainly located in the basalis gland of human endometria (Valentijn et al. 2013). Though mice quiescent epithelial label retaining cells were reported to be located mainly in the luminal epithelia and did not express the estrogen receptor (Chan and Gargett 2006), but their quiescence determined their difference with our proliferative EMT subset. The EMT epithelial cell subset in our study expressed higher level of CD44, a marker of the mouse endometrial epithelial progenitors located at both lumen and gland adjacent to the lumen that also express low level of hormone receptors (Janzen et al. 2013). However, the EMT epithelial cell subset in our study did not express SOX9, transcription factor reported to be vital for the regulation of endometrial epithelial stem/progenitor cells in the basal glands(Valentijn et al. 2013). Thus, our atlas provided a novel stem/progenitor cell subtype.

Cell plasticity was involved in the tissue development, homeostasis maintenance and pathological conditions (Varga and Greten 2017). Cell plasticity was increasingly reported in injury and regeneration of various epithelial tissues (e.g. intestine(Tetteh et al. 2016), liver, pancreas(Tritschler et al. 2017), hair follicles(Merrell and Stanger 2016)) in vivo. Endometrial cells were also reported to be highly plastic(Bilyk et al. 2017).Evidences already exist that EMT and MET involved in the menstrual regeneration and embryo implantation in the uterus(Bilyk et al. 2017). Subpopulation of MET cells in the regeneration zone that express both epithelial marker cytokeratin and stromal cell marker vimentin was involved in the menstrual endometrial regeneration (Patterson et al. 2013). MET of endometrial cells would make them more susceptible to deeper penetration by the embryo as epithelial cell gradually lose polarity and tight junctions (Paria et al. 1999).

EMT is also shown to properties of endometrial epithelial stem/progenitor cells(Wu et al. 2017), which may explain the in vitro culturing of endometrial epithelial stem/progenitor cells from menstrual shedding debris since 10 years ago(Gargett, Schwab, and Deane 2016). EMT was also involved in the pathogenesis, during which, epithelial cells would lose polarity and acquire motility, migration and proliferation, which would play a role in the evolution of adenomyosis, endometriosis and even cancer when dysregulated or under certain microenvironment(Bilyk et al. 2017).

### Our atlas provided a dynamic connectivity map of the uterus with diverse communications in the upper functionalis layer of the endometria regulating the EMT program

The low expression of hormone receptor in the reported stem/progenitor cells of SSEA1+ cells (Gargett, Schwab, and Deane 2016), Epithelial label retaining cells(Chan and Gargett 2006), CD44 + endometrial epithelial progenitors (Janzen et al. 2013) and ALDH1A1+ epithelial cells during development(Wu et al. 2017) implied that paracrine signals from the surrounding microenvironment were responsible for the mediation of hormonal regulation, a crosstalk between stem/progenitor cells with the surrounding niche cells was vital during the development, regeneration and might pathology(Gargett, Schwab, and Deane 2016; Wu et al. 2017; Chan and Gargett 2006; Janzen et al. 2013). Our atlas provided a dynamic connectivity map among subsets of the uterus.

TGF-beta family (TGF-beta, BMP), FGF, IGF1, EGF, PDGF, WNT were reported to regulate EMT through EMT transcription factors (SNAIL, TWIST, ZEB family), and inflammatory cytokines and hypoxia were shown to promote EMT by cross-talk with EMT TFs through STAT and HIF TF(Lamouille, Xu, and Derynck 2014)s. In the utrus, EMT was shown to be induced by estrogen through EMT TFs, and abnormal hormone level, dysregulation of WNT signaling would contribute to adenomyosis(Chen et al. 2010). Excessive ROS stress, abnormal exprsssion of ECM and ECM related LOX family, Lipocalin2 induced EMT is highly correlated with endometriosis through increased migratory and invasiveness properties(Vargha et al. 2008). Cytokines and chemokine, growth factors like TGFbeta signaling, hypoxia and oxidative stress are shown to be driver of EMT in endometrial cancer.(Bilyk et al. 2017)

## Conclusion

Here, we reconstructed an atlas of the human uterus tissues with the most detailed cell diversity, a compartmentalized cell ecosystem and a dynamic connectivity map of the human uterus tissue so far. Our atlas discovered new knowledge on uterus biology, would provide insight in the regeneration of uterus and reference for the pathogenesis of uterus.

## Materials and methods

### Human uterus collection

The Human full-thickness uterus sample was collected from the normal portions of the uterus of leiomyoma patients from the First Affiliated Hospital, School of Medicine, Zhejiang University. Approval for utilizing the patient samples in this study was obtained from Ethics Committee of the First Affiliated Hospital, School of Medicine, Zhejiang University.

### Single cell suspension preparation

Single cell suspension was prepared according to previous study(Turco et al. 2017). Briefly, full-thickness uterus tissue was minced into small cubes with scissors, and digested with collagenase V (Sigma) in RPMI 1640 medium (Thermo Fisher Scientific) with gentle shaking every 20-30 min at 37°C for 2hours. The stromal cells and smooth muscles were collected by passing the digested supernatant through 70μm cell sieves (Corning). The epithelial cell pellets were backwashed and further digested with TrypLE (Thermo Fisher Scientific) at 37°C for 10min. The digested supernatant was passed through 70μm cell sieves again to get single epithelial cell suspension. Finally, the stromal cells, smooth muscles and epithelial cells were combined to get the uterus single cell suspension for further analysis.

### Single cell capture, pre-amplification and sequencing

Single cell capture, pre-amplification was conducted onto the GemCode instrument (10x Genomics) according to the manufactures’ instructions (Chromium™ Single Cell 3′ Reagent Kit v2). The generated library was sequenced on five lanes of an Illumina X10 platform.

### Bioinformatic analysis

The generated sequencing reads were aligned and analyzed using the Cell Ranger Pipeline (10x Genomics). We obtained 2735 cells with about 680k reads per cell with a median gene number per cell of 3,183. Single cell analysis was conducted using Seurat(Satija et al. 2015). Pseudo-space was reconstructed using Monocle (Trapnell et al. 2014). Connectivity map was constructed according to (Puram et al. 2017; Camp et al. 2017) using ligand-receptor dataset(Ramilowski et al. 2016). Gene ontology analysis was conducted using: http://geneontology.org.

### Histology and Immunostaining

The human uterus tissue was fixed in 4% (w/v) paraformaldehyde, and then dehydrated in an ethanol gradient, prior to embedment in paraffin and sectioning at 10μm thickness. Immunostaining were carried out as follows: The 10μm paraffin sections were rehydrated, antigen retrieved, rinsed three times with PBS, and treated with blocking solution (1% BSA) for 30 min, prior to incubation with primary antibodies at 4 °C overnight. The primary antibodies: rabbit anti-human antibodies against COL3A1 (Abcam, ab7778), ZEB1 (Proteintech Group, 21544-1-ap), KI67(Abcam, ab16667), NFkB (Cell Signaling TECHNOLOGY, 3987), γH2A.X(Cell Signaling TECHNOLOGY, 2577), mouse anti-human monoclonal antibodies against Pan-KRT (Abcam, ab7753), CD45(BD Biosciences, 555483), CD68(Abcam, ab955) were used to detect the expression of selected proteins within the human uterus. TUNEL assay kit was used to detect apoptotic cells (TEASEN, China) within the human uterus. Secondary antibody: goat anti-rabbit Alexa Fluor 488 (Invitrogen, A11008), donkey anti-mouse Alexa Fluor 488 (Invitrogen, A21202), goat anti-rabbit Alexa Fluor 546 (Invitrogen, A21430-f) and DAPI (Beyotime Institute of Biotechnology, China) were used to visualize the respective primary antibodies and the cell nuclei. All procedures were carried out according to the manufacturer’s instructions.

## ACKNOWLEDGEMENTS

This work was supported by the National High Technology Research and Development Program of China (2017YFA0104902), the National Natural Science Foundation of China (CN) (81270682, 81300454), the Key Scientific and Technological Innovation Team of Zhejiang Province (2013TD11).

## AUTHOR CONTRIBUTIONS

B.W.: acquisition of clinical sample, data, data analysis and interpretation, manuscript writing; Y.L.: acquisition of sample, sample processing; Y.S.L.: processing of sample for immunostaining; K.X.J., K.Z., C.R.A., Q.K.L. data analysis; L.G.: manuscript preparation; W.Z., J.H.H., J.H.Q. acquisition of clinical sample; H.O.: conception and design, manuscript writing; X.Z.: conception and design, manuscript writing.

## Supplementary figures

**Figure S1.**
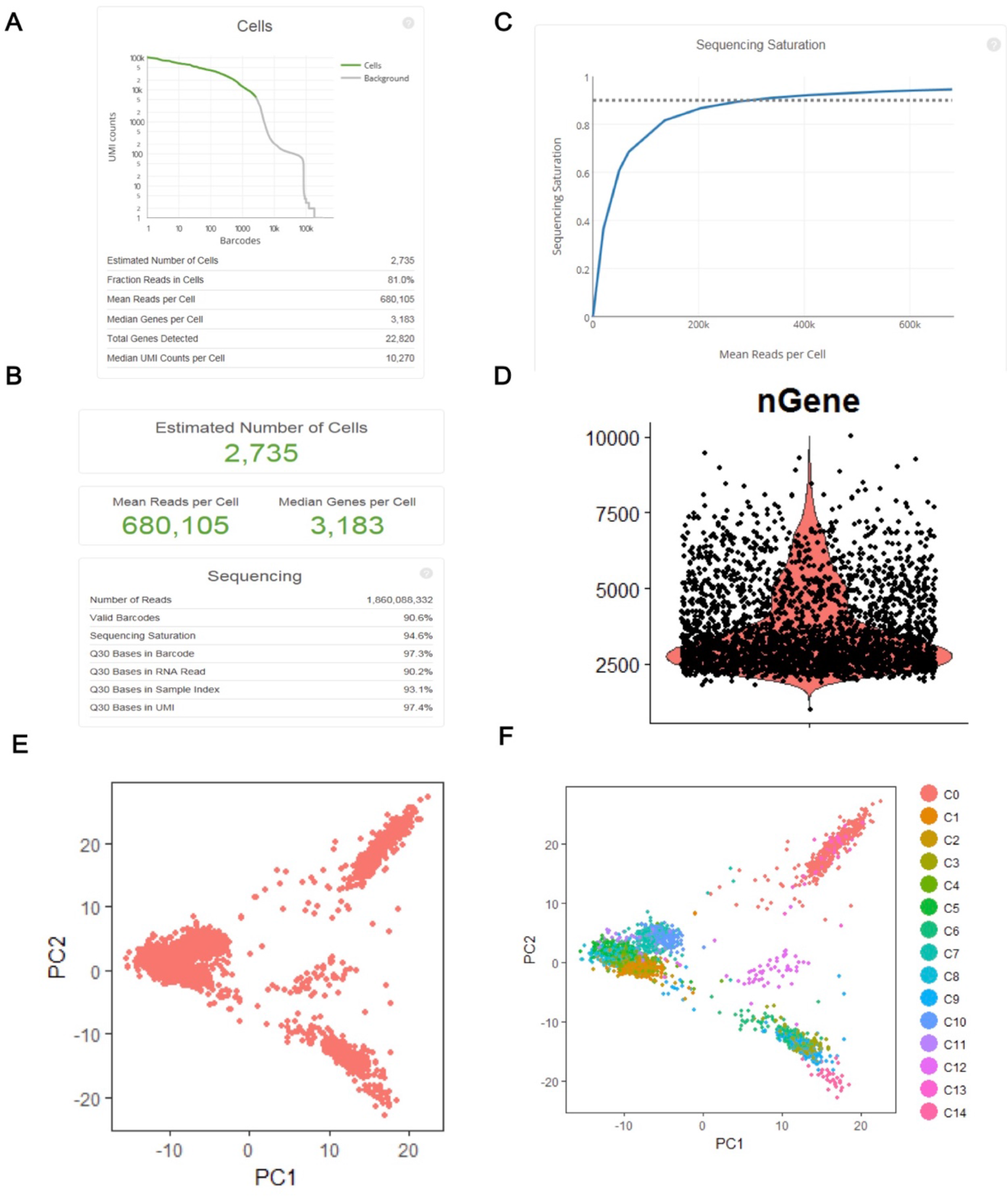
Quality control of the single cell RNA-seq. (A) We used the Cell Ranger Pipeline (10x Genomics) to analyze the unique molecular tagged (UMI) of the raw sequencing data. (B) We collected 2735 cells with high UMI counts of the human uterus. (C) We obtained saturated sequencing with about 680k reads per cell. (D) The median gene number detected per cell was about 3,183.(E)&(F) Unsupervised clustering based on principal components of the most variable expressed genes partitioned all the cells into 15 clusters, which was visualized with principal component analysis (PCA). Figure S1 was correlated with Figure 1.

**Figure S2.**
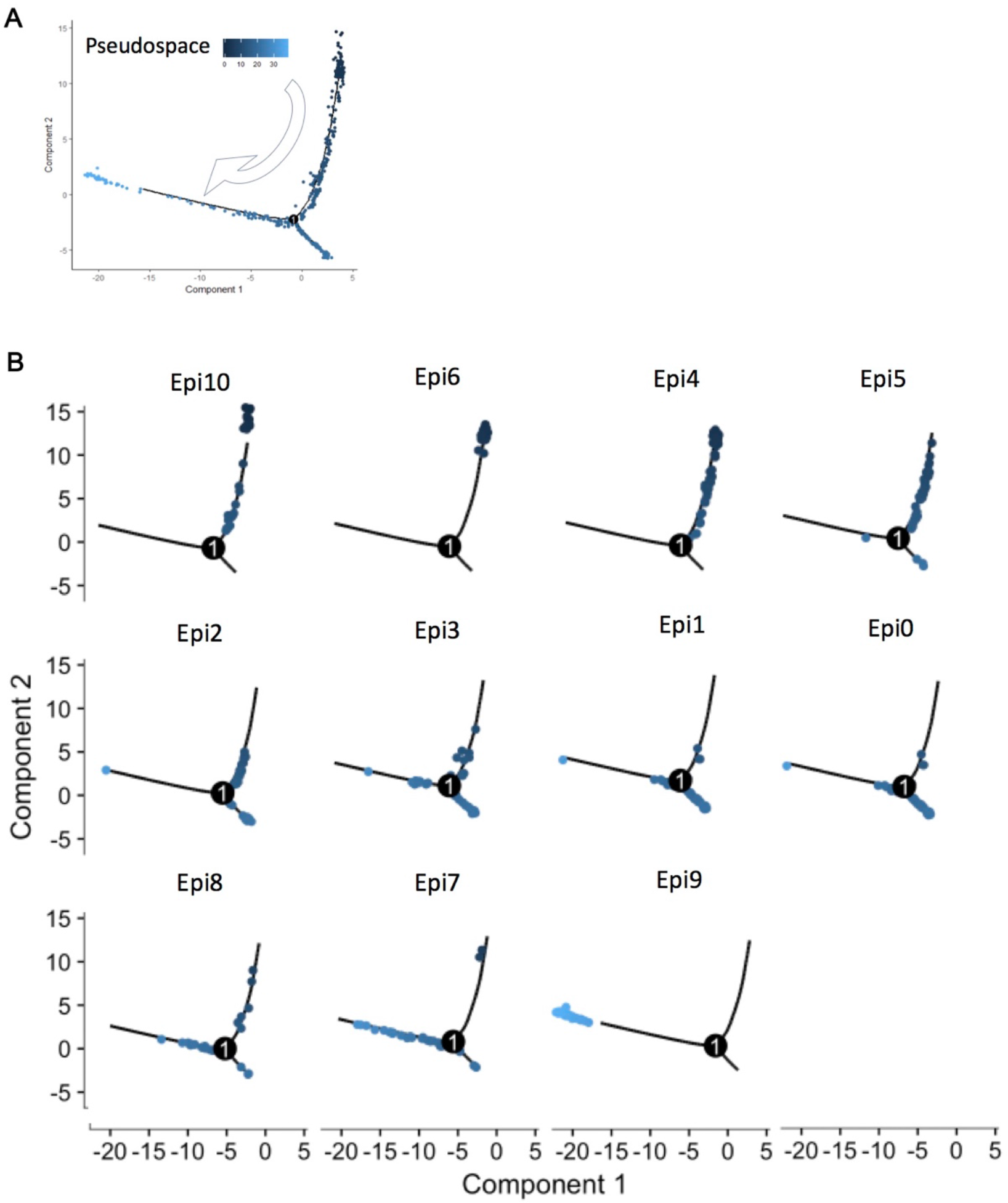
Pseudospace ordering of the uterus epithelia subsets. (A) Pseudospace ordering of all the epithelial subclusters. (B) Order of each epithelial sub-cluster in the pseudospace (Epi-Epi6-Epi4-Epi5-Epi2-Epi3-Epi1-Epi0-Epi8-Epi7-Epi9). Figure S2 was correlated with Figure 2.

**Figure S3.**
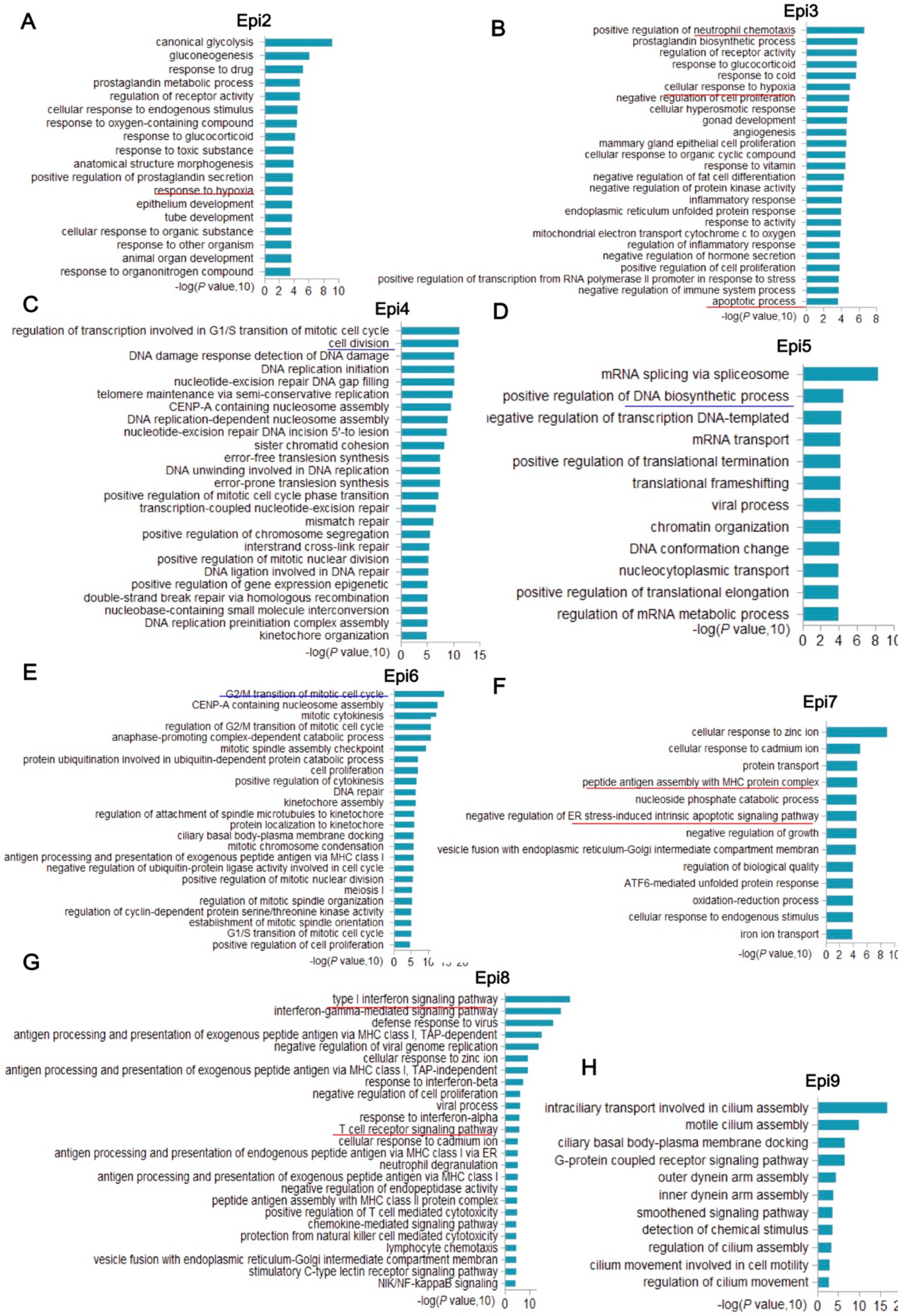
Gene ontology analysis of uterus epithelia subsets. **(A)** Gene ontology analysis of the highly expressed genes in the epithelial sub-cluster Epi2. **(B)** Gene ontology analysis of the highly expressed genes in the epithelial sub-cluster Epi3. **(C)** Gene ontology analysis of the highly expressed genes in the epithelial sub-cluster Epi4. **(D)** Gene ontology analysis of the highly expressed genes in the epithelial sub-cluster Epi5. **(E)** Gene ontology analysis of the highly expressed genes in the epithelial sub-cluster Epi6. **(F)** Gene ontology analysis of the highly expressed genes in the epithelial sub-cluster Epi7. **(G)** Gene ontology analysis of the highly expressed genes in the epithelial sub-cluster Epi8. **(H)** Gene ontology analysis of the highly expressed genes in the epithelial sub-cluster Epi9. Figure S3 was correlated with Figure 2.

**Figure S4.**
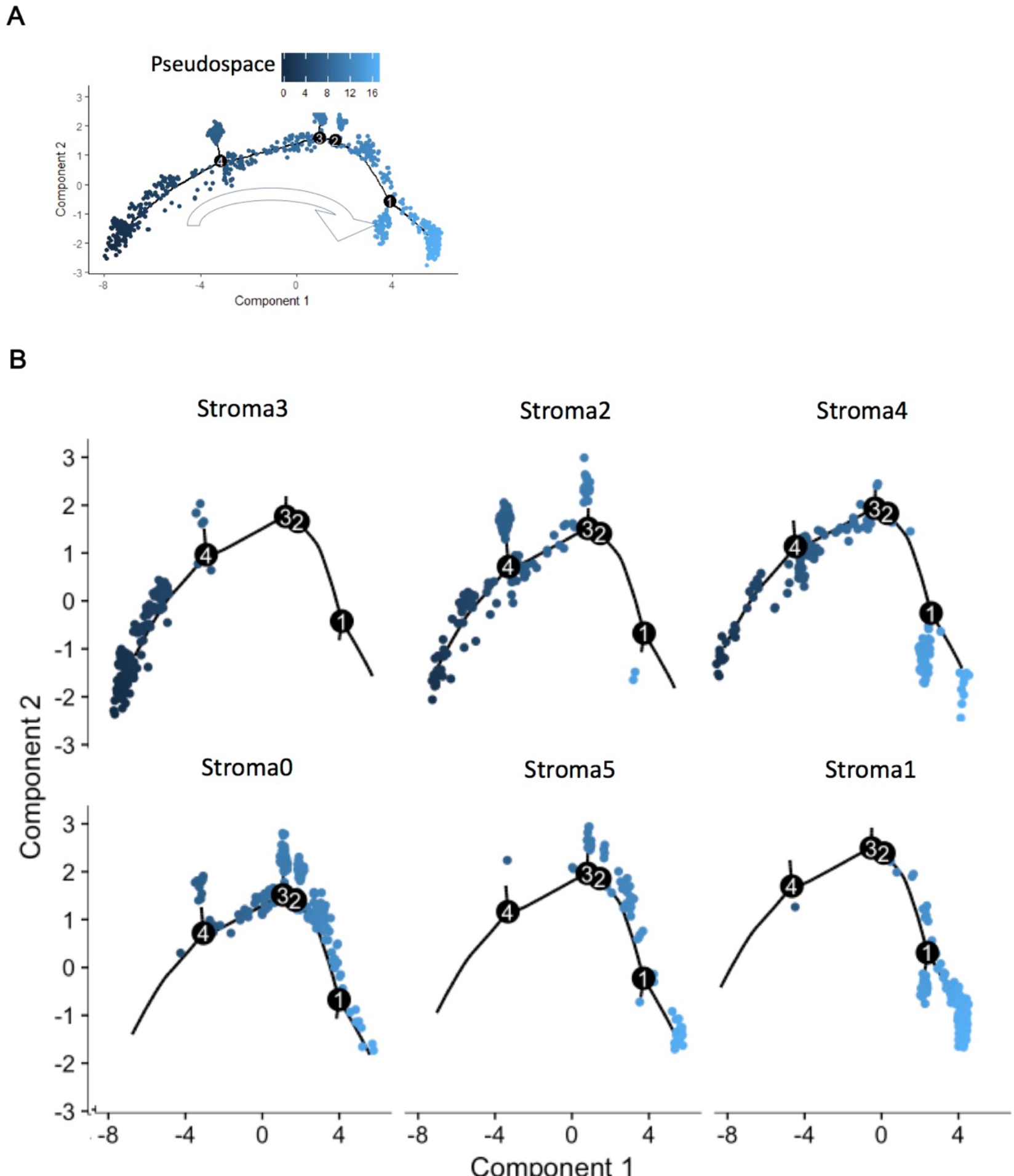
Pseudospace ordering of the uterus stroma subsets. (A) Pseudospace ordering of all the stromal subclusters. (B) Order of each stromal sub-cluster in the pseudospace (stroma3-stroma2-stroma4-stroma0-stroma5-stroma1). Figure S4 was correlated with Figure 3.

**Figure S5.**
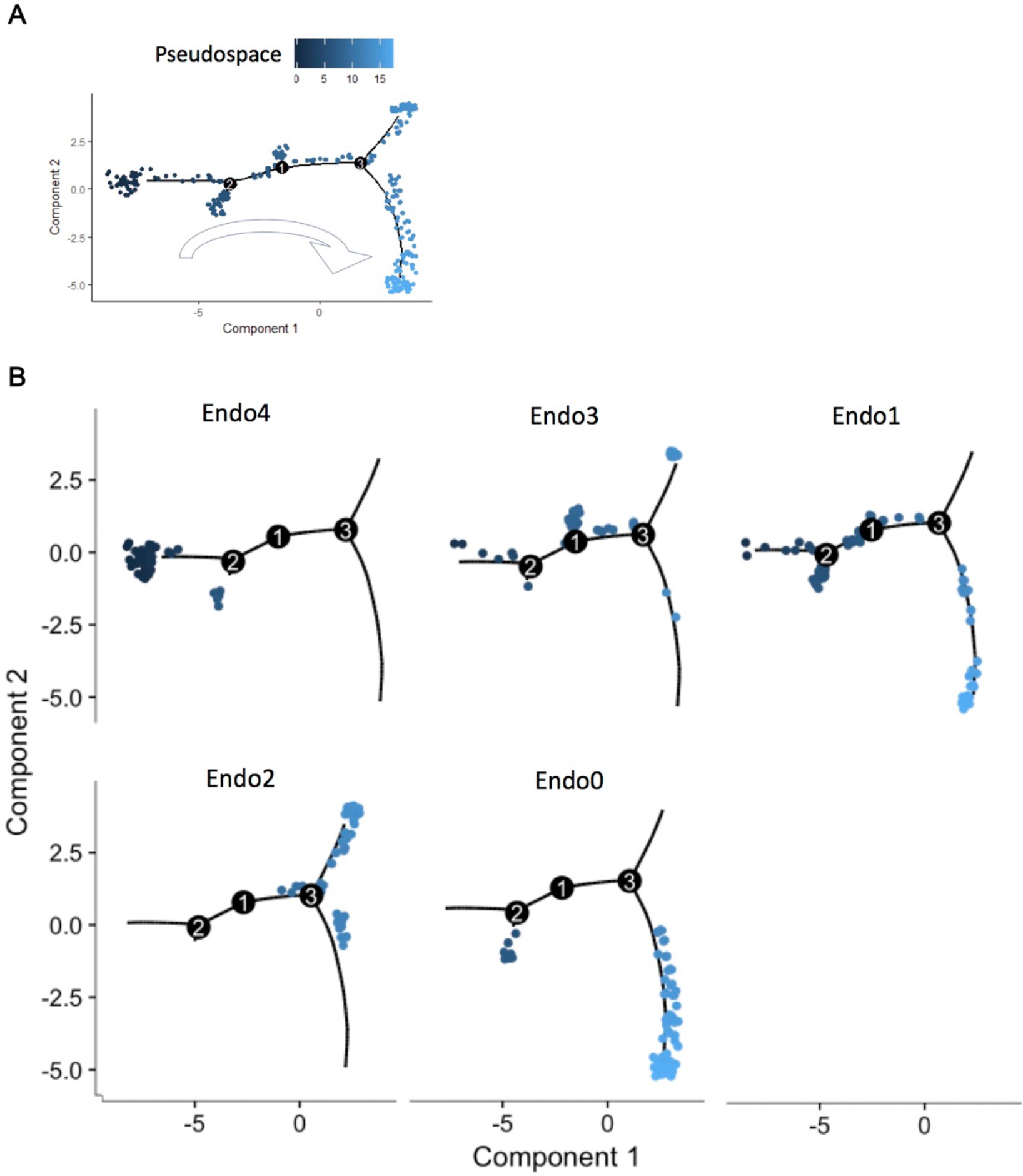
Pseudospace ordering of the uterus endothelial subsets. (A) Pseudospace ordering of all the endothelial subclusters. (B) Order of each endothelial sub-cluster in the pseudospace (Endo4-Endo3-Endo1-Endo2-Endo0). Figure S5 was correlated with Figure 4.

